# High-performance biodegradable membrane for point of need paper-based micro-scale microbial fuel cell analytical devices

**DOI:** 10.1101/351890

**Authors:** María Jesús González-Pabón, Federico Figueredo, Diana C. Martínez-Casillas, Eduardo Cortón

## Abstract

One limiting aspect to make microbial fuel cells (MFCs) a viable technology is to obtain low cost and environmentally sound materials for their components. In this work we synthesized membranes by a simple procedure involving low price and biodegradable materials such as poly (vinyl alcohol) (PVA), chitosan (CS) and PVA:CS, all cross-linked with sulfuric acid; they were compared to Nafion^®^, as our reference/control membrane. PVA:CS show lower oxygen permeability in comparison to Nafion^®^ membranes, a strong advantage in order to maintain anaerobic conditions in the anodic compartment of MFCs. Membranes were characterized in typical H-Type MFCs, and results show that PVA:CS membranes outperform Nafion^®^ 4 times (power production) while being 75 times more economic. Moreover, we design a paper-based micro-scale MFC, which was assayed as a toxicity biosensor; we obtained results in less than 20 min using 16 μL volume samples containing formaldehyde as a model toxicant. The PVA:CS membrane presented here can offer low environmental impact (materials, fabrication and disposal) and become a very interesting option for point of need single use disposable analytical devices.

## 1. Introduction

Microbial fuel cells (MFCs) are well known bio-electrochemical systems that can be used to understand how microorganisms manage redox process to sustain life. MFCs can also be used as a biotechnological tool, helpful in industrial and environmental areas. During the last decades, applied research involving MFCs technology was mainly focused in energy production (mostly electricity and hydrogen) and wastewater treatment processes, using both heterotrophic bacteria as biological catalysts and organic substrates as the fuel [1]. Later on, analytical applications of MFCs were developed, as biosensors for biochemical oxygen demand (BOD), lactate and acetate determination, toxicity and metabolic biosensors and even as life detectors, among others [2–4]. MFCs have emerged as a new type of analytical devices as they can be miniaturized and eventually able to produce enough power to become self-powered devices. In this way, MFC can be useful to measure relevant water quality parameters, whereas as a single use devices constructed with disposable and low cost materials [5–7], or well as a part of more sophisticated devices, as early warning toxicity systems are [8,9]. Moreover, a fuel cell transducer can be used not only for microbial-based biosensors, but also for enzymatic-based ones, where redox enzymes are typically employed [10].

There is a huge variety of MFCs designs, sizes and operation modes, depending on the intended use and materials availability. For example, large MFC reactors are required for wastewater treatment units with energy recovery; medium size systems can be enough to power up small electronic devices and very small paper-based and/or micro fabricated devices can be used as MFC-based analytical devices, given that only a small analytical signals (in the nA-mA range) are usually needed for calibration and quantification.

In a typical two chamber MFC, a PEM is used to separate the anodic and cathodic compartments, to avoid crossover of different dissolved substances, including gases. PEM membranes allow a selective transport of protons from the anode to the cathode whereas avoid the homogenization of the MFC compartments and prevents the oxygen transport into the anode chamber [11]. Nafion^®^ is a polymeric material with high mechanical strength, chemical stability, high electrical conductivity and selective permeability that is commonly used as PEM in MFCs [12–15]. However, Nafion^®^ has disadvantages like high cost (1767 USD/m^2^), especially when the goal is commercially available, single-use MFC-biosensors or other low cost MFC systems. Furthermore Nafion^®^ requires several activation steps, including a high temperature treatment [16]. Another Nafion^®^ drawback is it relatively high oxygen permeability, that allows oxygen leakage from cathode to anode [17]. Moreover, some of Nafion^®^ properties previously named (extraordinary chemical and thermal stability) become a problem when it needs to be disposed as waste; to be burned, the incinerator should have alkaline scrubber facilities to reduce hydrogen fluoride emissions to an acceptable amount, so recommended disposal is landfill, which is not the best option from an environmental point of view.

Several studies have explored the use of alternative membranes or separators in MFCs, such as cation or anion-exchange, glass fiber, osmosis or dynamic membranes; also earthenware, salt bridges and other materials or set-ups have been used in order to improve MFCs performance in some way and reduce their cost [18–20]. Composite materials formed for other polymers such as poly(ether sulfone) (PES), sulfonated poly(ether ether ketone) (SPEEK) and chitosan (CS) have been previously investigated as MFCs membranes with promising good results in terms of low cost, reduced biofouling and oxygen diffusion properties [21,22]. However, almost all of those studies were planned considering just the membrane performance, without evaluating the environmental impact of the materials synthesized. Ideally a good membrane for a disposable MFC not only has to perform properly, but also should be economic and biodegradable, in order to reduce the environmental impact and waste disposal cost.

Chitosan (CS) have been explored for membrane fabrication since it is an abundant and natural polymer [23–25]. Therefore, CS presents a number of intrinsic advantages such as biodegradability, biocompatibility, non-toxicity and good sorption properties [26]. Moreover, free amine and hydroxyl groups on the CS’s backbone are potential reactive sites that allow further CS modifications [27]. Cross-linking is the most widely used modification made for CS membranes construction; when CS is treated with sulfuric acid, the main chains of CS became ionically cross-linked, where the hydrogel turn relatively stable given the coulombic interaction between the amino groups in chitosan and sulfate ions. Conductivity and water uptake can increase with the sulfonation [28, 29]. Furthermore, CS can be blended with either hydrophilic or hydrophobic polymers (*e.g.* PVA) to enhance mechanical and thermal stability [30–32].

Point of Need (PON) devices allow a rapid *in-situ* determination of relevant parameters, mostly related to water quality (pH, conductivity, toxicity, single analyte determination, etc). In recent years, practical PON devices have been introduced by using cellulose filter paper and their close sub-products [33, 34]. These materials can be used for the development MFC biosensors used for toxicity and metabolisms detection [5, 6, 35, 36]. MFCs based biosensors to be used as PON devices need to be constructed with low cost materials (electrodes, membranes, etc), taking into account fabrication and waste disposal methods; all related in some way with the environmental impact of the designed product. Durability and mechanical stability of the materials become a minor concern, given that the fabricated device can be preserved (dehydrated) during the storage time and be functional to operate just enough time to finish the analysis, usually hours or less.

This work involves the synthesis of three economic and environmentally friendly (biodegradable) alternative membranes for MFCs devices, based on PVA, CS and PVA:CS. Relevant properties such as morphology, water uptake, conductivity, thickness, oxygen mass transfer and MFC-performance were assayed for all membranes prepared here and compared with a Nafion^®^ commercial membrane, used as a reference or control. After MFC performance was assayed in a classical two-compartment H-type cell, the best membrane (PVA:CS) was further incorporated into a disposable paper-based micro-scale MFC biosensor constructed with paper anode/cathode chambers. As a proof of concept it was assayed as a water toxicity biosensor, showing a fast response time (about 10 min) to 0.1% formaldehyde solution.

## 2. Materials and Methods

### 2.1 Reagents and Nafion^®^ commercial membranes

Poly (vinyl alcohol), degree of polymerization ~1600, degree of hydrolysis 97.5-99.5 mol % and chitosan (poly-(1,4)-β-N-acetyl-D-glucosamme) low molecular weight with around 50% deacetylation degree were supplied by Sigma Aldrich. Analytical grade acetic and sulfuric acids were obtained from the same company. Dextrose anhydrous, sodium sulphite, methylene blue, NaCl, KCl, K_2_HPO_4_ and formaldehyde 37% were also used. All the reagents were used without further purification and all the solutions were prepared with Milli-Q water. Nafion^®^ 117 membrane was obtained from DuPont Co. (Wilmington, DE, USA) and used after a recommended activation procedure, as detailed later in Section 2.3.

### 2.2 Membrane synthesis

Three types of membranes were synthesized by using solution casting and solvent evaporation technique, based on previously reported methodologies used to prepare membranes for different systems [28, 37–40]. After the synthesis procedure, all membranes were washed with Milli-Q water and stored at room temperature in a Falcon tube containing Milli-Q water. In brief, the procedure for the synthesis of each membrane is described below.

#### 2.2.1 CS membrane

Aqueous solution of CS (2 % w/v) was prepared by dissolving 1 g of chitosan in 50 mL of acetic acid aqueous solution (2 % v/v). The solution was stirred at 1000 rpm for 12 h at room temperature. After the complete dissolution of chitosan, the solution was filtered and stored at 4 °C for 24 h. Thereafter, 20 g of chitosan solution was casted on a glass Petri dish and left to dry for 24 h at room temperature, followed by a dehydration step for 6 h at 60 °C. The dried membrane was neutralized in NaOH 2M for 5 min and washed with abundant Milli-Q water. Then, the membrane was cross-linked by immersion in H_2_SO_4_ 0.5 M for 24 h at room temperature.

#### 2.2.2 PVA membrane

They were synthesized as follows: 5 g of PVA were added at 45 mL of Milli-Q water (10 % w/v aqueous solution of PVA) and hydrated for 24 h. Then, PVA was dissolved under stirring (500 rpm) at 80 °C for 2 h. Thereafter, 20 g of the homogenous PVA solution was casted on glass Petri dish and dried at 60 °C for 6 h. The dried membrane was dipped in H_2_O_2_ at 10 % v/v for 1 h, washed and then cross-linked by immersion in H_2_SO_4_ (10 % v/v) for 12 h.

#### 2.2.3 PVA:CS membrane

The prepared aqueous solutions of CS (Section 2.2.1) and PVA (Section 2.2.2) were mixed in a 1:1 proportion and stirred at 500 rpm for 2 h. After a complete mix, the solution was stored at 4 °C for 24 h. Then, 20 g of the resulting PVA:CS solution was casted on a glass Petri dish. It was left for 24 h at room temperature, followed by a dehydration process for 6 h in an oven at 60 °C. The membrane obtained was neutralized in NaOH 2M for 5 min and washed with abundant Milli-Q water. The crosslinking was performed by immersion in H_2_SO_4_ 0.5 M for 24 h at room temperature.

### 2.3 Membrane activations procedure

The Nafion^®^ 117 membrane was activated by successive immersions steps during 1 h (all warmed to 80 °C). First we use H_2_O_2_ (3 % v/v), second Milli-Q water, then H_2_SO_4_ 2 M, and finally Milli-Q water, as a procedure usually used in MFC area. The activation procedure improves the membrane performance, by increasing Nafion^®^ hydration and conductivity [13]. The three membranes synthesized in this work (CS, PVA and PVA:CS) were used without any activation procedure.

### 2.4 Surface topography study

Morphology of the membranes was observed with a field emission scanning electron microscope (FE-SEM Carl Zeiss NTS SUPRA 40, USA). The samples were dehydrated by immersion in alcohol solutions of 25, 50 and 100 % v/v followed by sputter-coating with a thin layer of gold (20 nm) using a current of 30 mA for 30 s.

### 2.5 Water uptake capacity

The water uptake capacity was determined by measuring membrane weight changes during the hydration process. The membranes were first dried in an oven at 30 °C for 15 h and then weighed (*W*_*dry*_). Once dried, membranes were immersed in Milli-Q water for an initial period of 1 min and after that, membranes were wiped with tissue paper and immediately weighed (*W*_*wet*_). This operation was repeated several times. Finally, the membranes were immersed in Milli-Q water and maintained at room temperature for 24 h. This measurement was conducted in triplicate. The water uptake (*W)* was calculated with the following equation [41, 42].

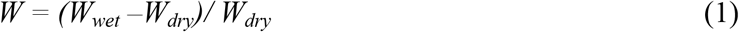

We express sometimes our results as % of water uptake (*W* × 100), for comparative proposes.

### 2.6 Conductivity determination

Four-point probe electrochemical impedance technique was used to determine proton conductivity of the synthesized and Nafion^®^ 117 membranes. The analysis was made scanning a frequency range between 10^−1^ and 10^6^ Hz at open circuit potential with an amplitude of 5 mV, using a commercial potentiostat (Interface 1000, Gamry, PA, USA).

A four electrode cell was constructed in the laboratory (Fig 1), by using Teflon blocks and Pd wires, according to previously reported work [28, 43]. When a fixed AC current flows between two outer electrodes, the conductance of the membrane can be calculated from the AC potential difference measured between the two inner electrodes. Thereby contact resistance, lead resistance and lead inductance do not interfere with the signal that is being measured [28].

**Fig 1.**
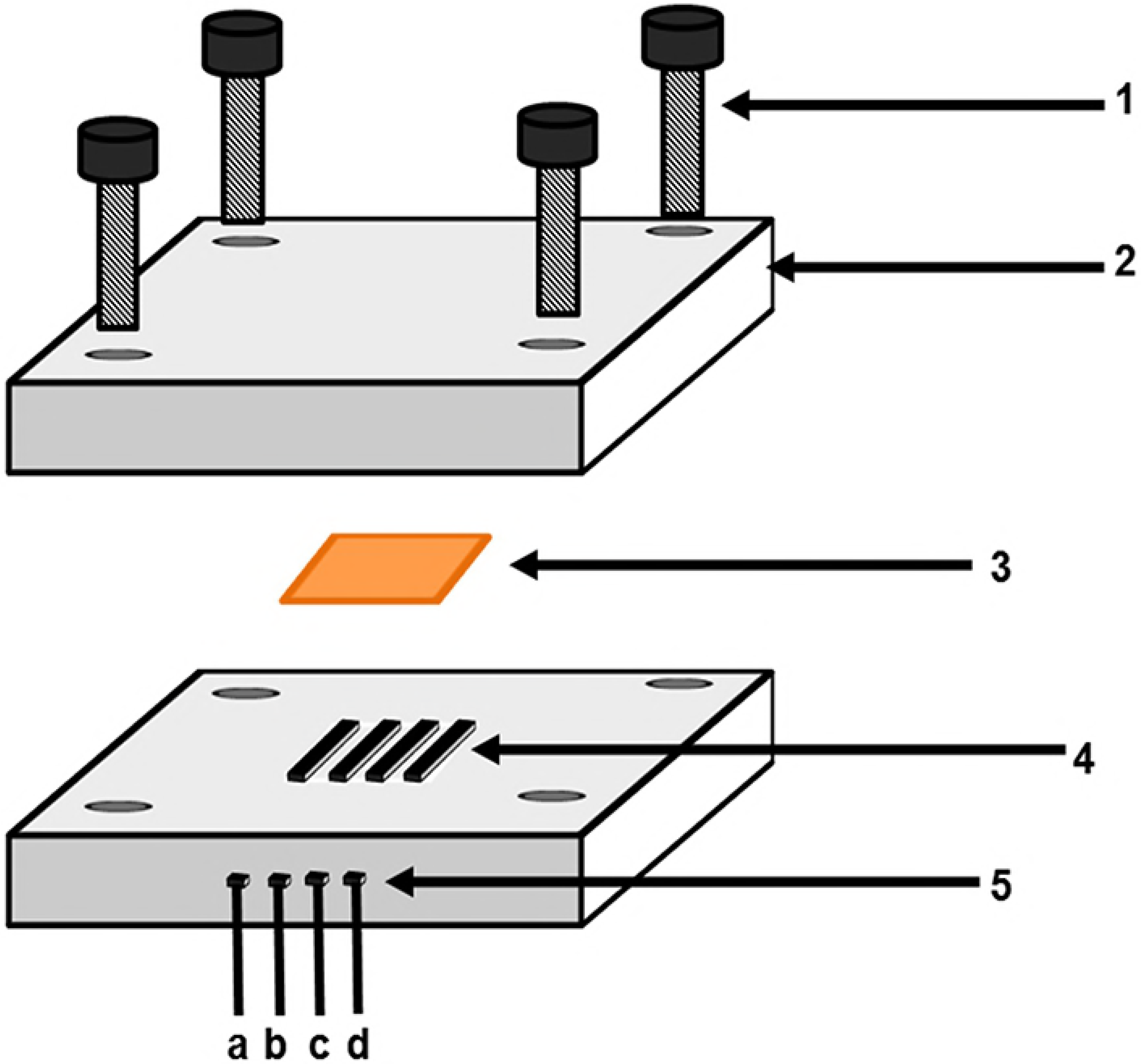
Conductivity cell and set-up. Membrane samples were measured by means of four-point electrochemical impedance spectroscopy. 1. Screws, 2.Teflon block, 3. Membrane sample, 4. Electrodes, 5. Electrodes contacts. **a**, **d** current carrying electrodes contacts. **b**, **c** potential-sensing electrodes contacts.

The analysis was performed at room temperature under 100 % RH (relative humidity), achieved by immersing the membranes in Milli-Q water before each measurement. In order to evaluate the reproducibility, each analysis was repeated three times. Gamry Echem Analyst software was used to simulate equivalent circuits and data tuning to extract the ohmic or bulk resistance of the membrane. The conductivity was calculated with the following equation:

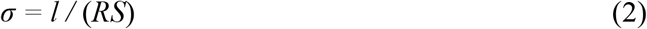

Where *σ* is conductivity (S/cm), *l* is the distance between the electrodes (cm), *R* is the ohmic resistance (Ω) of the membrane sample and *S* is the cross-sectional area of the membrane (cm^2^) [28].

### 2.7 Dissolved oxygen crossover determination

Oxygen transport was measured with an oxygen probe (oxygen meter Model D0-5510 by Lutron Electronic Enterprise Co., Ltd., Taipei, Taiwan). A small diffusion cell was used, made by replacing the original oxygen diffusion membrane of the oxygen electrode (as provided by the company) by the fabricated membranes (or Nafion^®^) to be assayed. Before each measurement, we proceeded to equilibrate the receptor (electrode chamber) and donor chambers, by bubbling N_2_ until a stable reading, close to 0 mg/L of dissolved oxygen (DO), was obtained (approximately 30 min). After that, the N_2_ stream was stopped and air was bubbled at the donor chamber. DO was monitored in the receptor chamber. The oxygen transfer coefficient 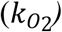 was determined using the mass balance equation:

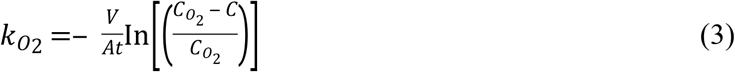

Where *V* is the receptor chamber volume (50 μL), *A* is the membrane cross-sectional area, 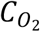 is the saturated oxygen concentration in the donor chamber, *C* is the DO concentration in the receptor chambers at time *t* [42, 44, 45]. A schematic representation of the used setup is presented (Fig A in S1 File). The oxygen diffusion coefficient (*Do*, cm^2^/s) was calculated replacing the membrane thickness (*L*_*t*_) in the follow equation:

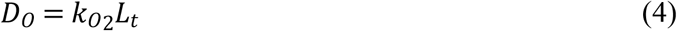

### 2.8 H-type MFC architecture and operation

The performance of membranes (1.3 cm^2^) was evaluated by polarization studies, by placing each membrane as separator of an H-Type MFC; anodic and cathodic compartments were of 16 cm^3^ each. All membranes used were disposed after each experiment (5 h of operation approximately). Toray paper was used as anode and cathode electrodes. The MFC was sterilized with 30% v/v H_2_O_2_ and 70% v/v ethanol for 15 minutes for each step. The cathode compartment was filled with 16 mL a solution containing potassium ferricyanide (50 mM, in order to avoid cathodic limitations) [2] and phosphate buffered (0.1 M, pH 6.2), whereas the anode compartment was filled with LB medium containing *E. coli* (OD = 1) and methylene blue (100 μM). It is worth noting that the aforementioned conditions (externally added mediator at the anode and ferricyanide cathode) are not sustainable when producing energy is intended; nevertheless they can perform perfectly as regents when an analytical goal is pursued (*i.e.*, MFCs-based biosensor). Before MFC measurements, the anode compartment was bubbled with N_2_ during 5 min to reach anoxic conditions. MFCs were maintained in a thermostatic chamber at 30° C during experiments. Potential measurements were done after 1 h at open circuit (OCV) and recorded with a data acquisition board (NI-USB 6210, National Instruments, USA) connected to a personal computer. Current (I, ampere) was calculated as I = E/R (5), where R was the external circuit resistor. Power (P, watt) was calculated as P = IE (6).

These values were normalized by using the electrode geometrical area (4 cm^2^ when H-type MFC was assayed), to obtain the current (*j*, A/m^2^) and power (*p*, W/m^2^) densities. Polarization and power curves (E vs. *j* and *p* vs. *j*, respectively) were obtained by applying different resistors to the circuit as an external load, ranging from 1 MΩ to 100 Ω. Resistors were connected sequentially, beginning from OC and then 1MΩ, to lower values, during 20 min each to allow stabilization. All H-type MFC experiments were carry out by duplicate.

### 2.9 Micro-scale, disposable MFC biosensor

The membrane that showed the best performance at H-type MFC experiments (Section 2.8) was also examined in a paper-based micro-scale MFCs. The membrane was used to separate anodic and cathodic reservoirs, made of filter paper (Whatman N° 1). Toray paper electrodes were used, details about the design and construction are presented at the supplementary information (Fig B in S1 File). The volume of each compartment was chosen to be 1000-folds smaller than the H-Type MFC used (Section 2.8), of 16 μL. The cathodic compartment solution was the same previously used (Section 2.8), whereas the anode compartment was filled with a solution containing *E. coli* (1.0 × 10^9^ CFU/mL) in a minimal medium constituted by phosphate buffer (0.1 M, pH 6.2), glucose (20 g/L), sodium sulfite (0.1 g/L) and methylene blue (100 μM). As a proof of concept, the paper-based micro-scale MFC was used as an acute toxicity biosensor; in such experiments formaldehyde (0.1%) was also added to the anolyte solution. MFC potential was continuously monitored by using a 100 KΩ load, and converted to current by using Ohm law (Eq. 5). MFCs were operated with toxic or control samples during 1 h and all the experiments were done at least by duplicate.

## 3. Results and Discussion

### 3.1 Surface topography study

FE-SEM pictures of CS membranes reveals a rough surface and porous structure, with pores of different diameters and heterogeneous distribution as can be seen in Fig 2a indicated by open arrows and in supplementary information (Fig C in S1 File). On the other hand, PVA and PVA/CS membranes show a non-porous structure, with a mostly smooth surface (Fig 2b and Fig 2c, respectively). PVA:CS membrane present features mainly of PVA, as the smooth surface and the presence of crystals, which can be seen in Fig 2c, indicated with solid arrows. Surface topology is an important factor that determines the fouling tendency of membranes, as well other properties. Rough surfaces foul more easily than smooth surfaces, due to the increase of surface area [22]. Hence, PVA and PVA:CS membranes will probably suffer less fouling than CS membrane.

**Fig 2.**
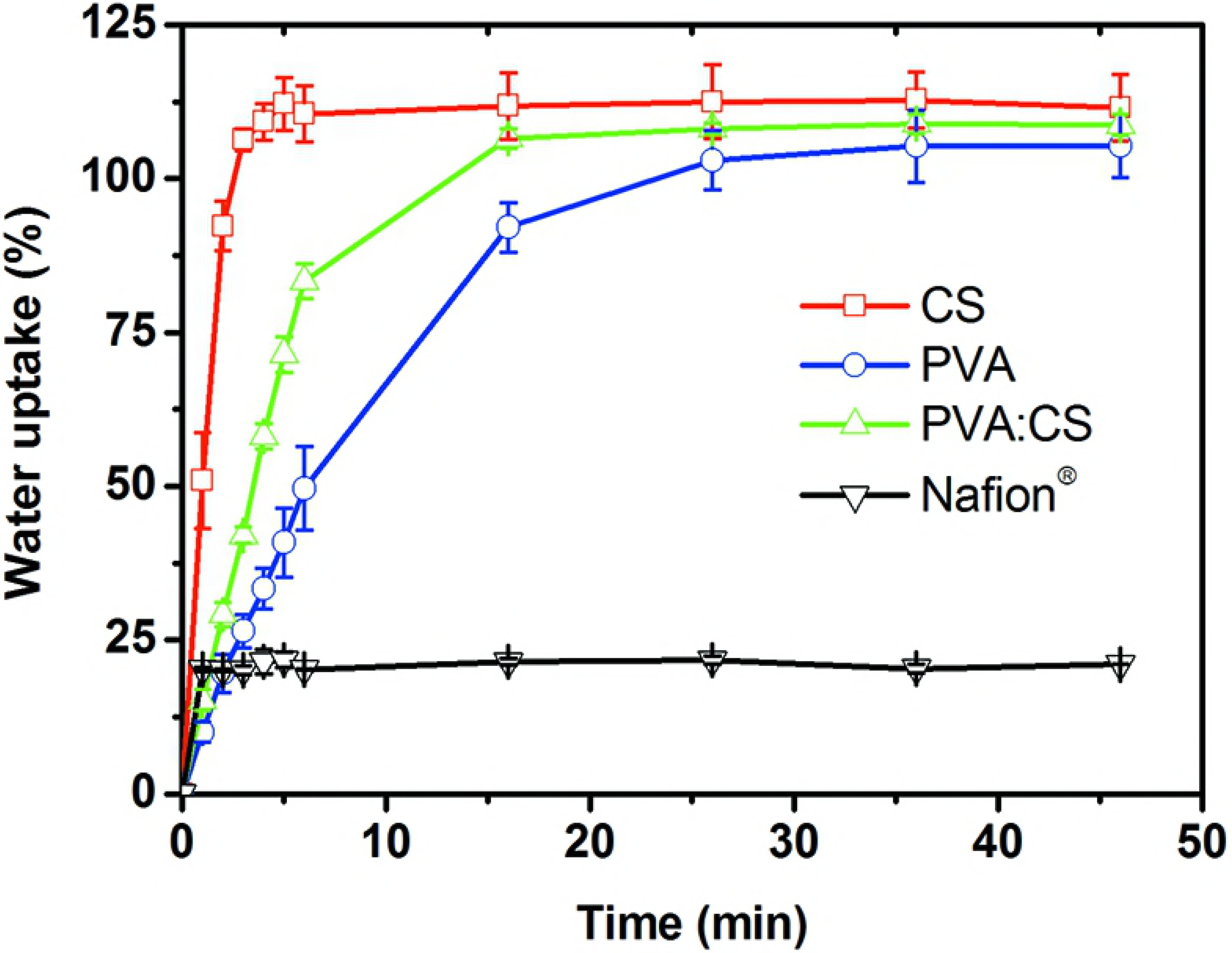
FE-SEM micrograph of the synthesized membranes. (a) CS, (b) PVA and (c) PVA:CS membranes. Open arrows indicates the pores, and solid arrows the PVA crystals.

### 3.2 Water uptake capacity

All membranes were quickly hydrated as is shown in Fig 3. All of them, including Nafion^®^ got full hydrated in less than 20 min. Water uptake was similar after 20 min and 24 hours for all membranes tested. Nafion^®^ water uptake value obtained was (23.32 ± 0.77) %, whereas synthesized membranes achieved more than 100% of their dry weight. Water uptake values obtained were (111.47 ± 3.28) % for CS, (105.18 ± 4.86) % for PVA and (108.73 ± 1.72) % for PVA:CS.

**Fig 3.**
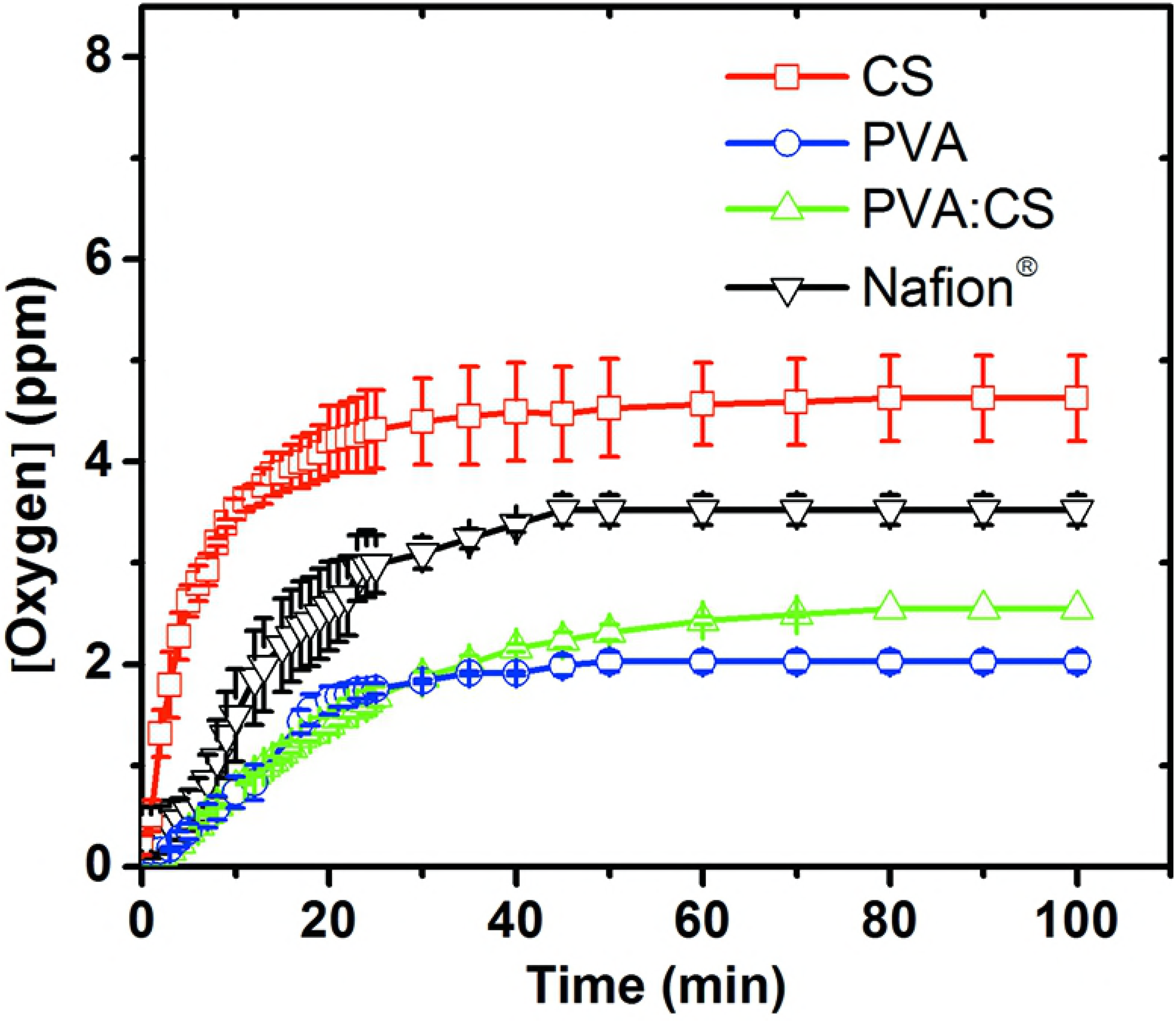
Kinetics of membrane hydration process. Water uptake of CS (□), PVA (○), PVA:CS (Δ), and Nafion^®^ (▽) membranes. Standard deviations are presented at all data points.

The higher water uptake shown by the synthesized membranes (when compared with Nafion^®^) could be a great advantage when considering hydrated proton transport in water, which can increase considerably the proton mobility [46]. If higher proton transport takes place, an enhanced MFC performance is expected.

### 3.3 Membrane conductivity

Membrane conductivity is an important aspect when parameters such as voltage and current of a MFC are analyzed. The mechanisms to describe the proton transfer (usually the main charge transporter) across the membranes are related to the ‘*Grotthus* mechanism’, where protons flow from one proton carrier to other, as-NH_2_,-NH_3_^+^ or -SO_3_H, which dissociate H_+_ and form hydrogen bonds. On the other side, there is a second mechanism named the ‘vehicle mechanism’, where protons are combined with water molecules to produce hydronium ions (*e.g.* H_3_O^+^, H_5_O_2_^+^, and H_9_O_4_^+^) that can migrate through a stream of water [41]. In this study four types of membrane were measured CS, PVA, PVA:CS and Nafion^®^, the last named is widely used in MFCs setups, allowing us sound comparisons against our in-house synthesized membranes. The proton conductivities obtained were 11.9, 3.9, 11.3 and 81.0 mS/cm for CS, PVA, PVA:CS and Nafion^®^, respectively. The CS and PVA:CS membranes displayed higher conductivity values than PVA membranes. The obtained conductivity values were in the same order of magnitude according to previously reported work (Table A in S1 File).

### 3.4 Membrane dissolved oxygen crossover

The oxygen diffusion through the membrane is a key factor in MFCs operation, since anoxic conditions are necessary in the anodic chambers to avoid competition between the electrode and oxygen as electron acceptors. Also, oxygen can be toxic for bacterial species that have obligate anaerobic metabolism [17]. CS membrane shows the higher oxygen mass transfer 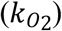, of 6.67 × 10^−4^ cm/s (Fig 4). This can be related to it porous structure, as revealed in SEM pictures (Fig 2a), which can allow easy oxygen permeation; this is an undesirable characteristic for their use in MFCs. PVA and PVA:CS membranes were very similar in their behavior 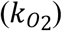, their oxygen permeability was approximately 4 times lower than CS membranes (Fig 4). These are the two membranes with better characteristic with respect to this parameter, to be used as MFC membranes. Nafion^®^ shows an intermediate oxygen diffusion coefficient (*Do*), lower than CS and higher than PVA and PVA:CS. Table 1 summarize the oxygen mass transfer coefficient 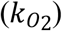 and oxygen diffusion coefficient obtained (*Do*) for all the membranes studied here.

**Fig 4.**
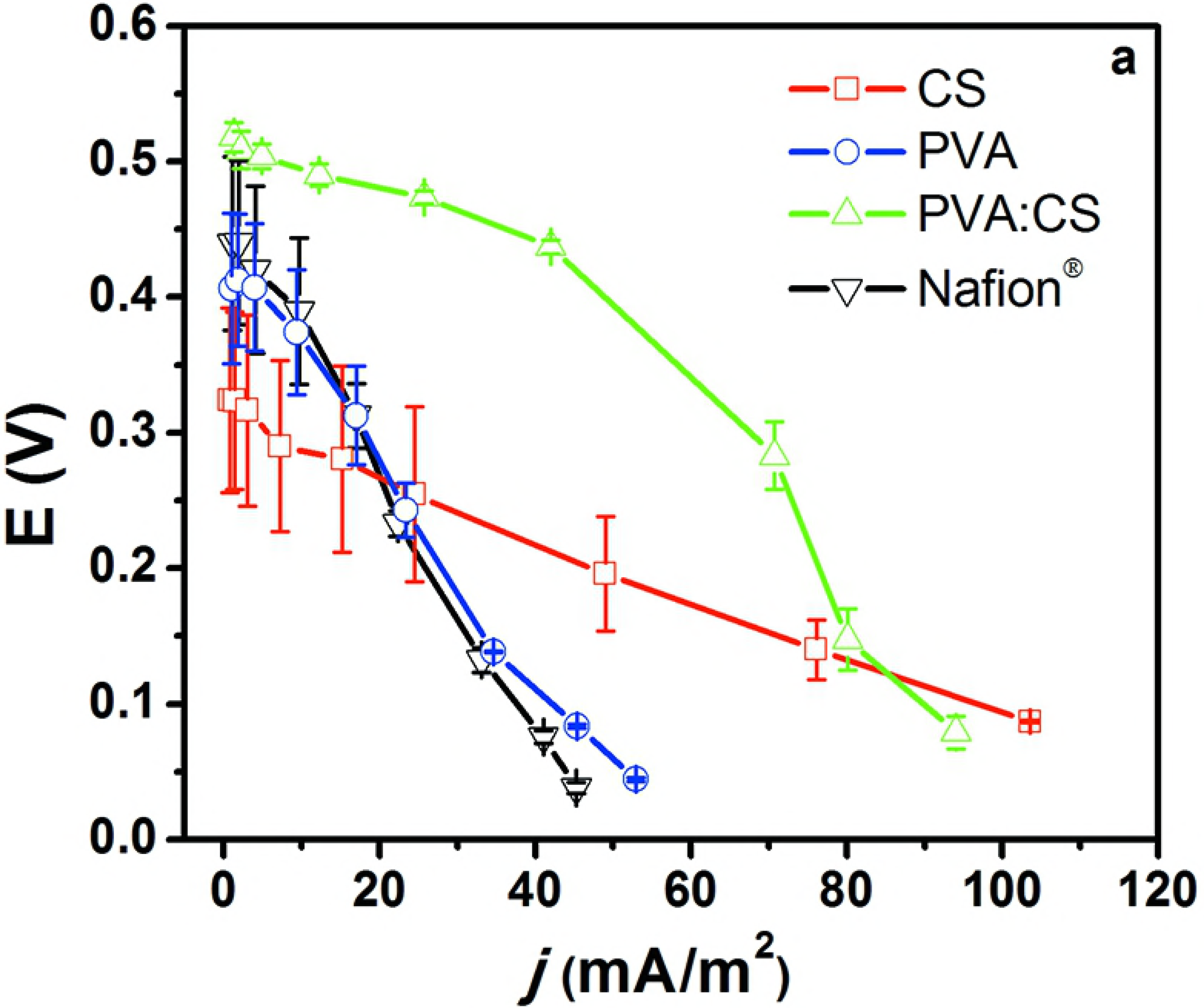
Oxygen diffusion across membranes. (□) CS, (○) PVA, (Δ) PVA:CS, and (▽) Nafion^®^ (reference membrane). Standard deviations are presented for all data points.

**Table 1.** 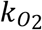 and *D*_*O*_ for CS, PVA, PVA:CS and Nafion^®^ membranes. 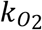 and *Do* were calculated from Eq. 3 and Eq. 4, respectively.

| Membrane | $10^4 k_{O_2}$ (cm/s)<br>(RSD%) | $10^6 D_o$ (cm<sup>2</sup>/s)<br>(RSD%) |
| --- | --- | --- |
| CS | 6.67 (9.8) | 6.67 (9.8) |
| PVA | 1.70 (5.6) | 2.04 (5.6) |
| PVA:CS | 1.50 (1.5) | 1.99 (1.5) |
| Nafion<sup>®</sup> | 2.71 (10.8) | 4.07 (10.8) |

### 3.5 MFCs performance in H-type cells

MFC performance depends to a great extent on each particular configuration, architecture, materials, microorganism and operation mode chosen. In order to evaluate the membrane’s effects under practical operational conditions, all components used in this study were identical for the MFCs tested, so the unique variable was the membrane. From the comparison between the three membranes synthesized here and Nafion^®^ 117, we made our results stronger and comparable with respect to other published work. First, the MFC’s OCV was measured for 1 hour and after 30 min, we obtained a stable OCV value. Then, the performance of each MFC was studied after the polarization experiments were made; adjusting the external load on the circuit, and measuring the stabilized potential (at 20 min). Polarization curves are presented in Fig. 5. Among the membranes tested in the MFCs, MFCs with the Nafion^®^ and PVA membranes showed the lowest maximum power (*p*_max_), of about 5.6 ± 0.1 and 5.7 ± 0.9 mW/m^2^ respectively. On the other hand, CS containing membranes showed higher values of current (at their maximum power values, *J*_max_) of 70.8 ± 6.3 and 76.1 ± 11.9 mA/m^2^ for PVA:CS and CS, respectively, and a *p*_max_ of 20.8 ± 2.9 and 11.5 ± 2.7 mW/m^2^ for PVA:CS and CS, respectively (Fig 5).

It is well known that membranes affect the internal resistance (R_int_) and therefore the global performance in the MFC system [42]. Nonetheless, the CS membrane shows the lowest internal resistance, and achieves the second-best power density. This result could be attributed to the high oxygen permeability of this membrane (Table 1) due to its porous structure. When oxygen is transported from the cathodic compartment into the anodic compartment, the oxygen acts as final electron acceptor instead of the electrode. As consequence, there are energy losses, and in general the MFC’s efficiency is low. This demonstrates that the oxygen permeability is a key factor determining the MFC performance [17]. Moreover, the MFCs assembled with the CS membranes showed the highest *J*_max_ values, which also made them an alternative membrane to be used in single use disposable MFC devices. However, the heterogeneous porous structure (Fig 2a) seems to produce low reproducibility and high deviations as can be seen in Fig 5. On the other hand, the PVA:CS membranes showed great characteristics related to the oxygen permeability, conductivity, and power density in MFC assays; these results indicate that this membrane has the potential to be employed as alternative to Nafion^®^ at least in disposable, low power MFC systems. The relevant data obtained here are summarized in the Table 2, which shows that PVA:CS membranes are fourfolds more efficient than Nafion^®^ 117 membranes, when *p*_max_ is compared. Low cost membranes as designed here are a perfect match for the construction of MFCs paper-based devices, to be used in PON applications and easy to dispose by burning, sometimes the only available disposal method in low-income countries.

**Fig 5.**
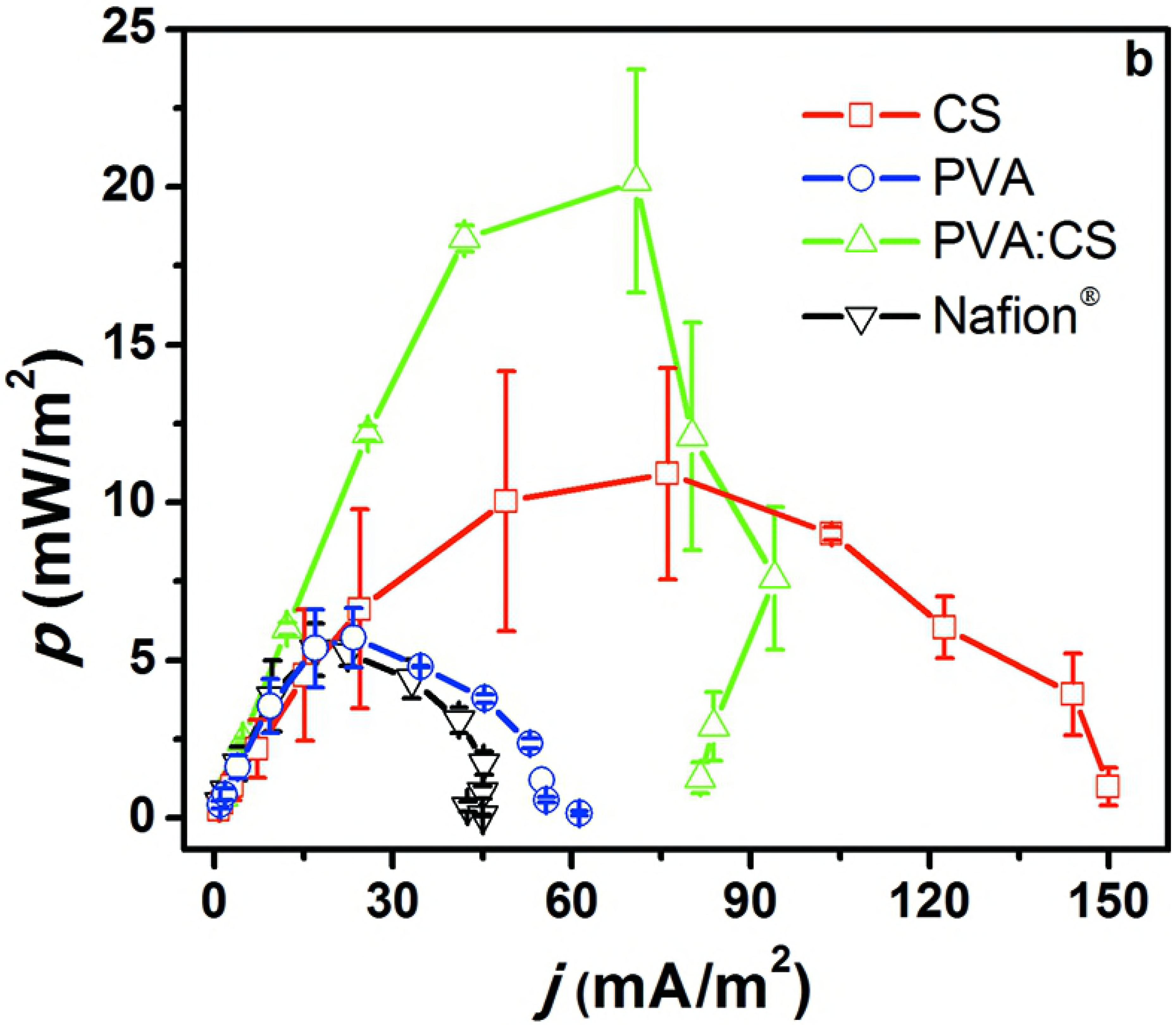
H-Type MFC performance. **a.** Polarization curves and **b.** Power density curves of (□) CS, (○) PVA, (Δ) PVA:CS, and (▽) Nafion^®^ membranes. Standard deviations are presented at all data points.

**Table 2.**
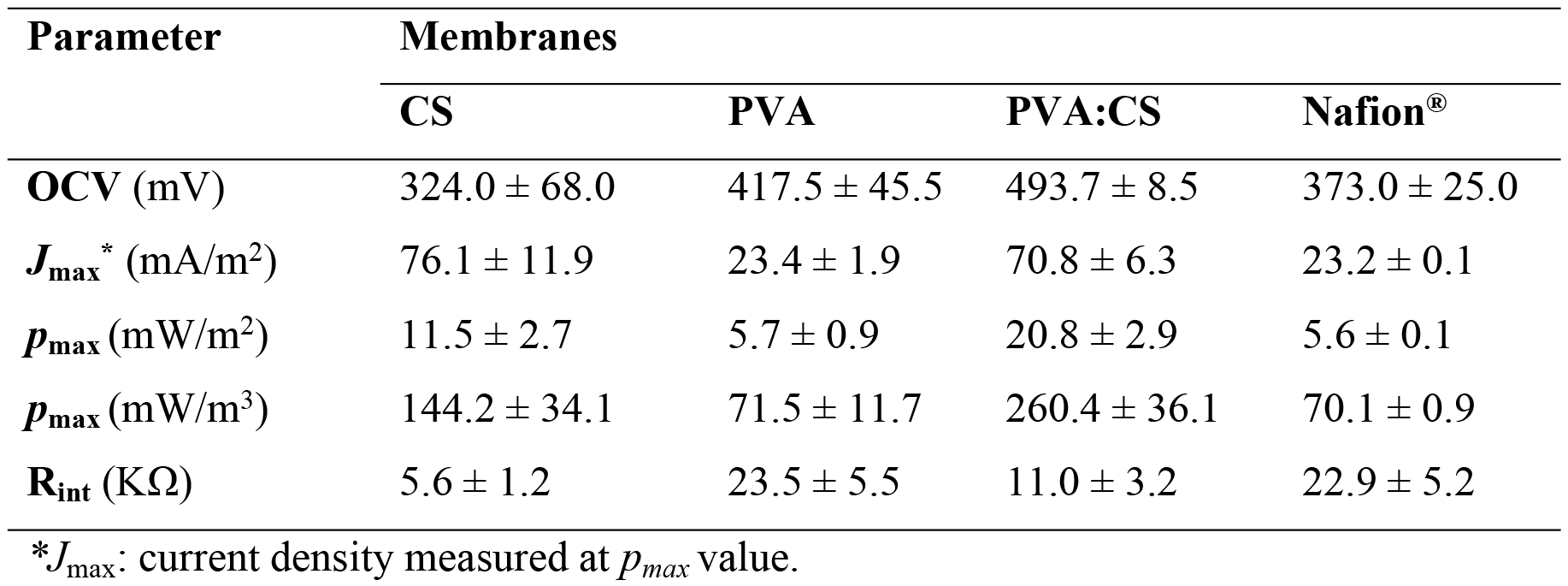
Relevant characteristic of different membranes assayed in an H-type MFC set-up.

### 3.6 Micro-scale and disposable MFC biosensor

As a proof of concept, we demonstrate the construction and use of the PVA:CS membrane as a fundamental part of a disposable paper-based micro-scale MFC biosensor used for toxicity determination. This kind of technology could replace or complement those commercial toxicity bioassays as Microtox in low resources countries as PON devices. We expose the microbial population present in the anodic compartment to formaldehyde (0.1%), which affected the current produced by the MFC, showing a decrease of about 64% ± 3% when compared with the control, without any toxic compound. The response was evident just 10 min after toxic exposure. The use of PVA:CS membranes in a simple and economic paper-based micro-scale MFC open a myriad of new analytical possibilities, where PON analysis are required, and simple disposal procedures (as incineration) must be used.

**Fig 6.**
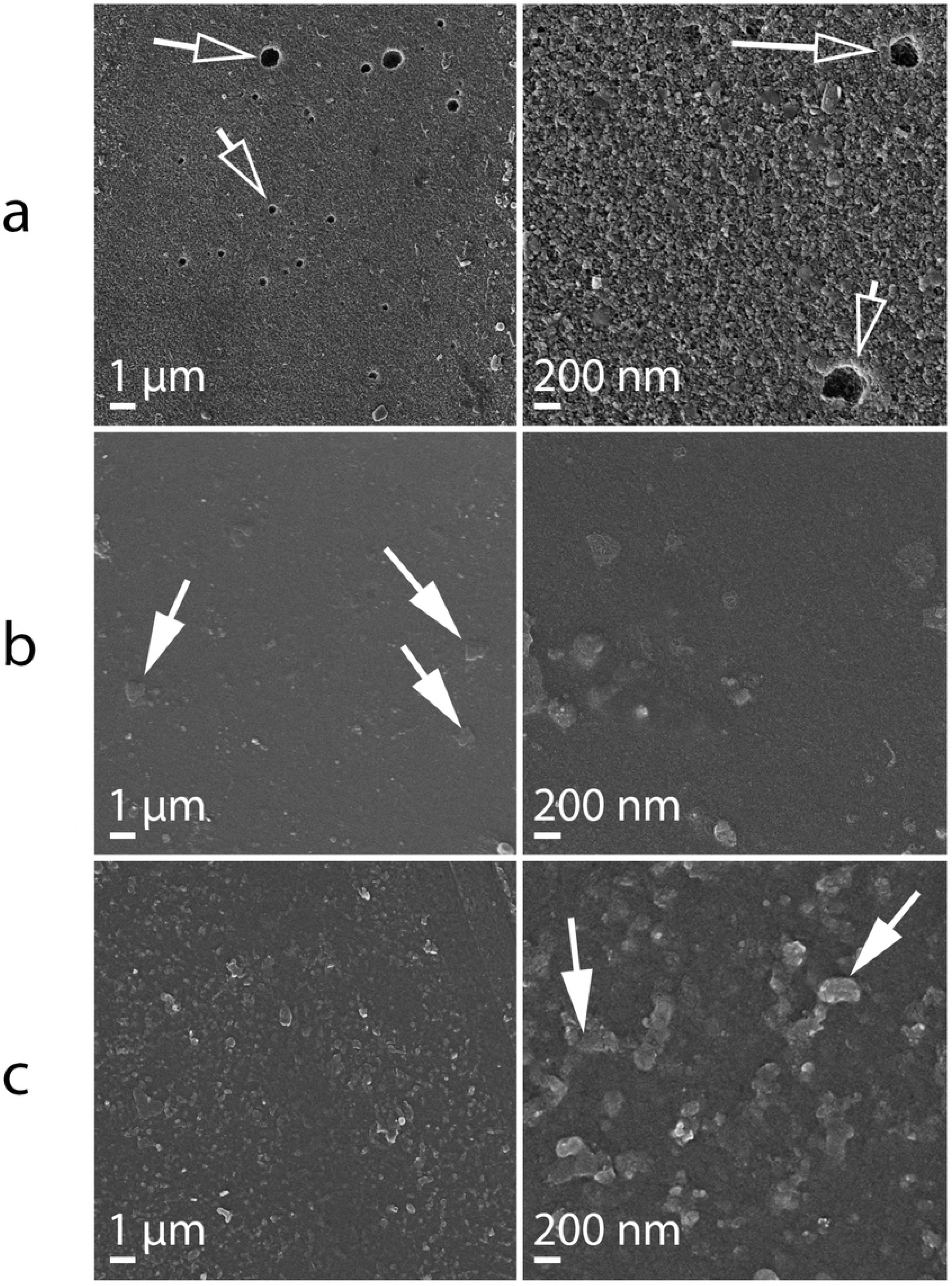
Paper-based micro-scale MFC as toxicity sensor. Current response of paper based MFC after the addition of a toxic sample containing 0.1% of formaldehyde compared to a control sample. Standard deviations are presented at all data points.

Schematic diagram of the paper-based micro-scale MFC device and polarization curve could be seen in supplementary information (Fig B in S1 File). The micro-scale MFC assayed here, proposed as simple PON analytical device shown several valuable analytical characteristics. OCV was relatively constant during the first 10 min of operation (dV/dt < 10%, data not shown), displaying short stabilization times, which is important for short time analysis, being a characteristic advantage of micro-scale MFC systems [47]. Moreover, after connecting the resistor, the systems becomes stable and remains stable during enough time gather accurate data (t = 10-30 min, Figure 6).

The micro-scale MFC device presented here, which does not rely on the typically used biofilm systems, allows a faster start-up, and so the delivery of the analytical answer. Moreover, as the device is made by low cost and biodegradable materials, durability and stability of such materials are expected to be modest, but a good choice when short term operation is part of the design. When very low volume systems are considered, evaporation can be a problem; still, systems as the proposed here could be eventually by useful for emergency power generation, by using large paper-based dehydrated devices [48].

### 3.7 Cost comparison between Nafion^®^ 117and synthesized membranes

Materials and reagents needed for the preparation of the best-performing membrane assayed in a MFC set-up (PVA:CS) are shown in Table 3.

**Table 3.** Cost of the reagents needed to make 1 m^2^ of PVA:CS membrane.

| Raw materials | Price, USD/kg | Amount, Kg | USD / m<sup>2</sup> |
| --- | --- | --- | --- |
| PVA | 5 | 0.20 | 1.0 |
| CS | 40 | 0.04 | 1.6 |
| NaOH | 5 | 0.80 | 4.0 |
| H<sub>2</sub>SO<sub>4</sub> | 5 | 0.45 | 2.25 |
| Total membrane |  |  | 8.85 |

We considered the values offered for bulk quantities, as found in Alibaba.com webpage. CS and PVA were of pharmaceutical and cosmetic grade, respectively. Manufactured Nafion^®^ membrane reach values about 2000 USD/m^2^ in the market [49]. Nafion^®^ resin was found at about 1845 USD/kg in the aforementioned webpage. 1 kg of resin would be enough to made approximately 2.8 m^2^ of membrane (360 g/m^2^). So, the material cost would be close to 659 USD/m^2^.

The Table 3 shows a calculated cost of around 8.9 USD/m^2^ for the PVA:CS membrane, meaning that PVA:CS copolymer is not only more efficient when used in a MFC, but also cost-effective, about 75 times cheaper. Furthermore PVA and CS are biodegradable materials. CS is a natural polymer and PVA, according to previous work, could be degraded by microorganisms like *Pseudomonas* sp., *Alcaligenes* sp., *Bacillus* sp. and *Phanerochaete crysosporium* [50]. Low cost and easy to dispose materials are fundamental aspects when a disposable PON device in on consideration.

### 3.8 Comparison with other low cost membranes

Table 4 shows a list of low cost membranes recently tested (in chronological order with the exception of Nafion^®^, in the first line). A relationship between the *p*_Max_ of different alternatives membranes reported and Nafion^®^ was calculated as can be seen in the following equation:

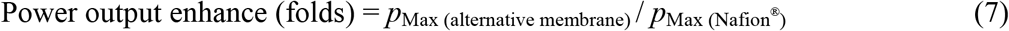

Where, *p*_Max(Altemative membrane)_ is the maximum power obtained with the proposed membrane and *p*_Max(Nafion^®^)_ is the maximum power obtained with Nafion^®^ membrane.

All values were obtained in the same experimental conditions and MFC set-up. MFC performance is contrasted with Nafion^®^ values present in each work (Table 4), necessary to made sound comparisons among different set-ups; some previously presented work where a new membrane is presented but not compared with Nafion^®^ were not included in Table 4.

**Table 4.**
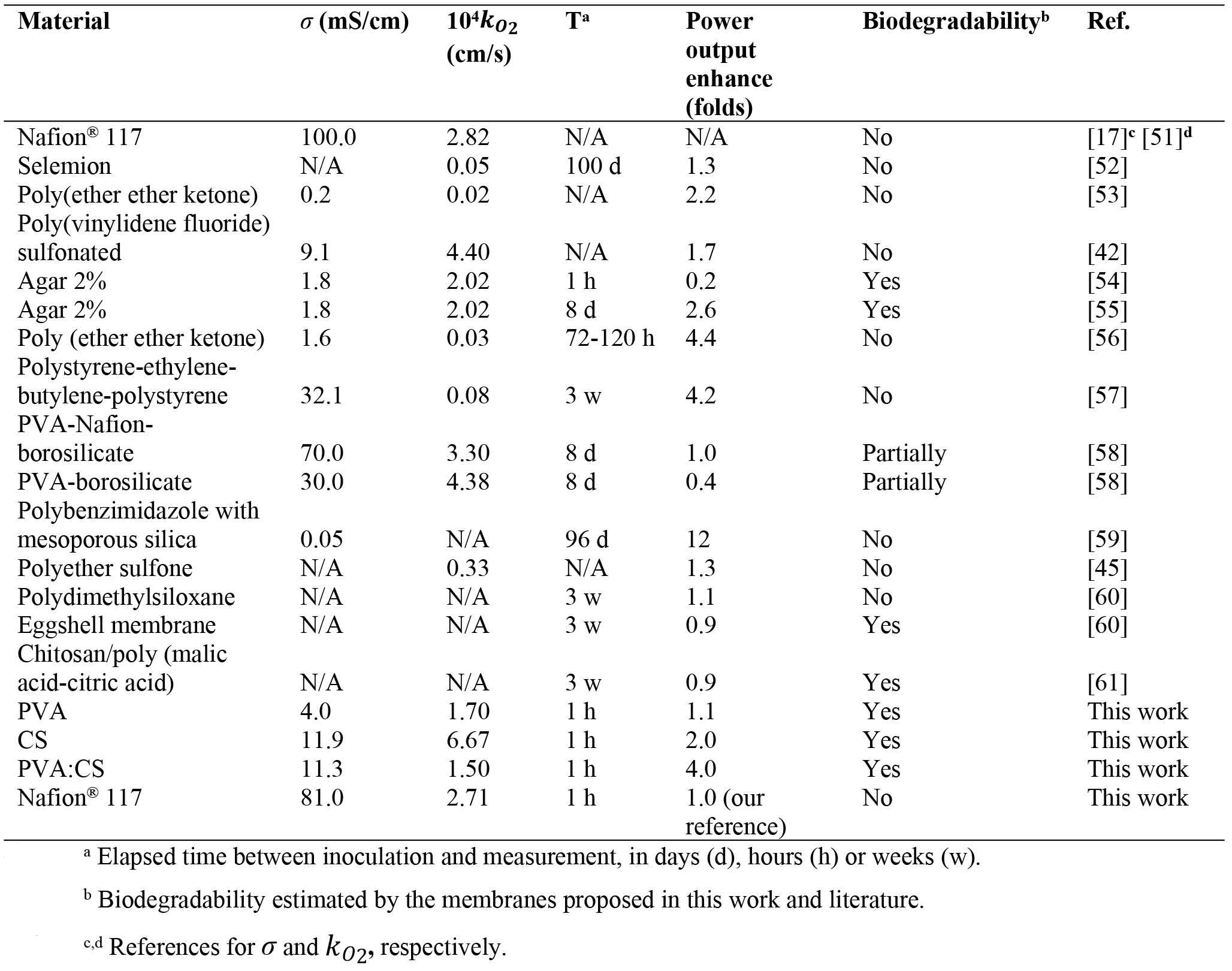
General features of low cost membranes in the literature and biodegradability comparisons. Power output enhace was compared to Nafion^®^, with respect with the values calculated at each reference.

## 4. Conclusions

Three types of membranes cross linked with sulphonic groups were investigated and characterized here by SEM and EIS, as well as their water uptake, oxygen diffusion and MFC performance. All our data was compared with Nafion^®^ 117, so the differences found here will be mostly related to the membrane characteristics and not by the set-up. From the three membranes synthesized, PVA:CS shows the best performance. One of the major advantages of PVA:CS membrane is that the oxygen permeability measured was lower than Nafion^®^117. Although, conductivity values of PVA:CS membrane was lower than Nafion^®^ 117, we found very good performance of the PVA:CS membrane when it was incorporated to an H-Type MFC and into a paper-based micro-scale MFC. This behavior could be related to the interactions of all the characteristics of each membrane type and strongly dependent on the use we made of them.

We described here a high performance, biodegradable and cost-effective PVA:CS membrane capable of outperforming Nafion^®^117 at least in the conditions we assayed, related with low-density power systems, as MFC. The maximum power achieved when used in a H-Type MFC were 4 times higher with PVA:CS, whereas the cost was about 75 times lower, which gives a superiority factor (efficiency times cost) of about 300 times when compared to Nafion^®^.

PVA:CS can overcome some relevant MFC bottlenecks, given its low cost. As a bonus, synthesized membranes can be prepared easily without the use of dangerous materials including organic solvents and can be easily disposable, given fast biodegradability and non-toxicity character. These are all optimal features for cost-effective, point of need paper-based disposable analytical systems, as paper MFC-based biosensors. As a proof of concept, the paper-based micro-scale MFC performs as expected to detect water containing a toxic substance, working as a fast (10 min) water quality sensor.

## Declaration of interest

The authors have no conflicts of interest to declare.

## Supporting Information

**S1 File. (Fig A)** Schematic representations of the setup for oxygen mass transport coefficient determination. **(Fig B)** a. Micro scale MFC construction diagram: materials, dimensions in mm and electrical connection. b. Paper-based micro-scale MFC performance. Polarization curves (square) and power density curves (circle). **(Fig C)** FE-SEM micrograph of the CS membrane. **(Table A)** Proton conductivity of the membranes prepared here compared with values published by others.

## Acknowledgements

We want to thank to MSc candidate Sebastian Cortón for editing our manuscript.

## Author Contributions

Conceived and designed the experiments: MJG, FF, DM, EC. Performed the experiments: MJG, DM, FF. Analyzed the data: MJG, DM, FF. Contributed reagents/materials/analysis tools: MJG, DM, FF. Wrote the paper: MJG, FF, EC. Writing - review & editing: MJG, EC. Funding: EC. Project administration: EC.

---

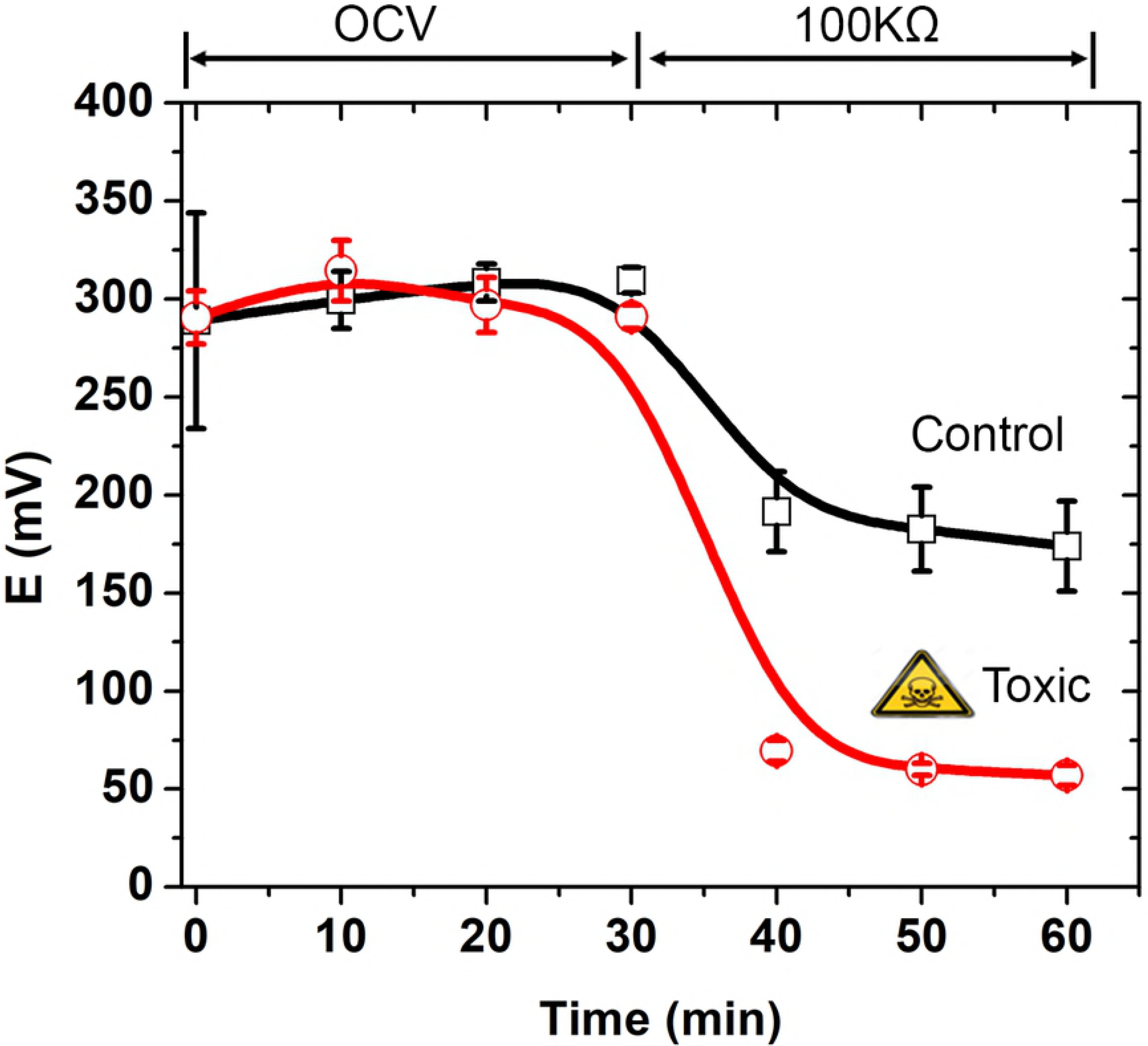

## References

[1] Pant D, Singh A, Van Bogaert G, Irving Olsen S, Singh Nigam P, Diels L, et al. Bioelectrochemical systems (BES) for sustainable energy production and product recovery from organic wastes and industrial wastewaters. RSC Adv. 2012; 2: 1248–1263. https://doi.org/10.1039/c1ra00839k.

[2] Abrevaya XC, Sacco NJ, Bonetto MC, Hilding-Ohlsson A, Corton E. Analytical applications of microbial fuel cells. Part I: biochemical oxygen demand. Biosens Bioelectron. 2015a; 63: 580–590. https://doi.org/10.1016/_j.bios.2014.04.034.

[3] Abrevaya XC, Sacco NJ, Bonetto MC, Hilding-Ohlsson A, Cortón E. Analytical applications of microbial fuel cells. Part II: toxicity, microbial activity and quantification, single analyte detection and other uses. Biosens Bioelectron. 2015b; 63: 591–601. https://doi.org/10.10167j.bios.2014.04.053.

[4] Figueredo F, Cortón E, Abrevaya XC. In situ search for extraterrestrial life: a microbial fuel cell-based sensor for the detection of photosynthetic metabolism. Astrobiology. 2015; 15: 717–727. https://doi.org/10.1089/ast.2015.1288.

[5] Choi, S. Microscale microbial fuel cells: advances and challenges. Biosens Bioelectron. 2015; 69, 8–25. https://doi.org/10.1016/_j.bios.2015.02.021.

[6] Choi G, Hassett DJ, Choi S. A paper-based microbial fuel cell array for rapid and high-throughput screening of electricity-producing bacteria. Analyst. 2015; 140(12): 4277–4283. https://doi.org/10.1039/C5AN00492F.

[7] Figueredo F, González-Pabón MJ, Cortón E. Low cost layer by layer construction of CNT/chitosan flexible paper-based electrodes: a versatile electrochemical platform for point of care and point of need testing. Electroanal. 2018; 30(3): 497–508. https://doi.org/10.1002/elan.201700782.

[8] Xu GH, Wang YK, Sheng GP, Mu Y, Yu HQ. An MFC-based online monitoring and alert system for activated sludge process. Sci Rep. 2014; 4: 6779. https://doi.org/10.1038/srep06779.

[9] Jia H, Yang G, Wang J, Ngo HH, Guo W, Zhang H, et al. Performance of a microbial fuel cell-based biosensor for online monitoring in an integrated system combining microbial fuel cell and up flow anaerobic sludge bed reactor. Bioresour Technol. 2016; 218:286–293. https://doi.org/10.1016/_j.biortech.2016.06.064.

[10] Bollella P, Fusco G, Stevar D, Gorton L, Ludwig R., Ma S, et al. A glucose/oxygen enzymatic fuel cell based on gold nanoparticles modified graphene screen-printed electrode. Proof-of-concept in human saliva. Sensor Actuat B-Chem. 2018; 256: 921–930. https://doi.org/10.1016Zj.snb.2017.10.025.

[11] Christgen B, Scott K, Dolfing J, Head IM, Curtis TP. An evaluation of the performance and economics of membranes and separators in single chamber microbial fuel cells treating domestic wastewater. PloS One. 2015. https://doi.org/10.1371/journal.pone.0136108.

[12] Unnikrishnan EK, Kumar SD, Maiti B. Permeation of inorganic anions through Nafion ionomer membrane. J Memb Sci. 1997; 137: 133–137. https://doi.org/10.1016/S0376-7388(97)00193-2.

[13] Mauritz KA, Moore RB. State of understanding of Nafion. Chem Rev. 2004; 104: 4535–4586. https://doi.org/10.1021/cr0207123.

[14] Stenina IA, Sistat P, Rebrov AI, Pourcelly G, Yaroslavtsev AB. Ion mobility in Nafion-117 membranes. Desalination. 2004; 170 49–57. https://doi.org/10.1016/j.desal.2004.02.092.

[15] Lehmani A, Turq P, Périé M, Périé J, Simonin JP. Ion transport in Nafion^®^ 117 membrane. J Electroanal Chem. 1997; 428: 81–89. https://doi.org/10.1016/S0022-0728(96)05060-7.

[16] Hernández-Flores G, Poggi-Varaldo HM, Solorza-Feria O, Ponce-Noyola MT, Romero-Castañón T, Rinderknecht-Seijas N, et al. Characteristics of a single chamber microbial fuel cell equipped with a low cost membrane. Int J Hydrogen Energ. 2015; 40: 17380–17387. https://doi.org/10.1016/j.ijhydene.2015.10.024.

[17] Chae KJ, Choi M, Ajayi FF, Park W, Chang IS, Kim IS. Mass transport through a proton exchange membrane (Nafion) in microbial fuel cells. Energ Fuels. 2008; 22: 169–176. https://doi.org/10.1021/ef700308u.

[18] Daud SM, Kim BH, Ghasemi M, Daud WRW. Separators used in microbial electrochemical technologies: current status and future prospects. Bioresour Technol. 2015; 195: 170–179. https://doi.org/10.1016/j.biortech.2015.06.105.

[19] Kadier A, Simayi Y, Abdeshahian P, Azman NF, Chandrasekhar K, Kalil MS. A comprehensive review of microbial electrolysis cells (MEC) reactor designs and configurations for sustainable hydrogen gas production. Alexandria Eng J. 2016; 55: 427–443. https://doi.org/10.1016/j.aej.2015.10.008.

[20] Li X, Liu G, Sun S, Ma F, Zhou S, Lee JK. Power generation in dual chamber microbial fuel cells using dynamic membranes as separators. Energ Convers and Manage. 2018; 165: 488–494. https://doi.org/10.1016/j.enconman.2018.03.074.

[21] Wu H, Fu Y, Guo C, Li Y, Jiang, N, Yin C. Electricity generation and removal performance of a microbial fuel cell using sulfonated poly (ether ether ketone) as proton exchange membrane to treat phenol/acetone wastewater. Bioresource Technol. 2018; 260: 130–134. https://doi.org/10.10167j.biortech.2018.03.133.

[22] Antolini E. Composite materials for polymer electrolyte membrane microbial fuel cells. Biosens Bioelectron. 2015; 69: 54–70. https://doi.org/10.1016Zj.bios.2015.02.013.

[23] Batista MKS, Pinto LF, Gomes CAR, Gomes P. Novel highly-soluble peptide-chitosan polymers: chemical synthesis and spectral characterization. Carbohyd Polym. 2006; 64: 299–305. https://doi.org/10.1016yj.carbpol.2005.11.040.

[24] Dashtimoghadam E, Hasani-Sadrabadi MM, Moaddel H. Structural modification of chitosan biopolymer as a novel polyelectrolyte membrane for green power generation. Polym Adv Technol. 2009; 21: 726–734. https://doi.org/10.1002/pat.1496.

[25] Kalaiselvimary J, Sundararajan M and Prabhu MR. Preparation and characterization of chitosan-based nanocomposite hybrid polymer electrolyte membranes for fuel cell application. Ionics. 2018; 1–17. https://doi.org/10.1007/s11581-018-2485-7

[26] Witt MA, Barra GMO, Bertolino JR, Pires ATN. Crosslinked chitosan/poly (vinyl alcohol) blends with proton conductivity characteristic. J Braz Chem Soc. 2010; 21: 1692–1698. http://dx.doi.org/10.1590/S0103-50532010000900014.

[27] Ye YS, Rick J, Hwang BJ. Water soluble polymers as proton exchange membranes for fuel cells. Polymers. 2012; 4: 913–963. https://doi.org/10.3390/polym4020913.

[28] Mukoma P, Jooste BR, Vosloo HCM. Synthesis and characterization of crosslinked chitosan membranes for application as alternative proton exchange membrane materials in fuel cells. J Power Sources. 2004; 136: 16–23. https://doi.org/10.1016/jjpowsour.2004.05.027.

[29] Ma J, Choudhury NA, Sahai Y, Buchheit RG. A high performance direct borohydride fuel cell employing cross-linked chitosan membrane. J Power Sources. 2011; 196: 8257–8264. https://doi.org/10.1016/jjpowsour.2011.06.009.

[30] Xu D, Hein S, Wang K. Chitosan membrane in separation applications. Mater Sci Technol. 2008; 24: 1076–1087. https://doi.org/10.1179/174328408X341762.

[31] Rhim J, Park H, Lee C, Jun J, Kim D, Lee Y. Crosslinked poly (vinyl alcohol) membranes containing sulfonic acid group: Proton and methanol transport through membranes. J Memb Sci. 2004; 238: 143–151. https://doi.org/10.1016/j.memsci.2004.03.030.

[32] Pivovar BS, Wang Y, Cussler EL. Pervaporation membranes in direct methanol fuel cells. J Memb Sci. 1999; 154: 155–162. https://doi.org/10.1016/S0376-7388(98)00264-6.

[33] Xia Y, Si J, Li Z. Fabrication techniques for microfluidic paper-based analytical devices and their applications for biological testing: a review. Biosens Bioelectron. 2016; 77: 774–789. https://doi.org/10.1016/j.bios.2015.10.032.

[34] Martinez AW, Phillips ST, Butte MJ, Whitesides GM. Patterned paper as a platform for inexpensive, low-volume, portable bioassays. Angew Chem Int. 2007; 46: 1318–1320. https://doi.org/10.1002/anie.200603817.

[35] Chouler J, Cruz-Izquierdo A, Rengaraj S, Scott JL, Di Lorenzo M. A screen-printed paper microbial fuel cell biosensor for detection of toxic compounds in water. Biosens Bioelectron. 2018; 102: 49–56. http://dx.doi.org/10.1016/j.bios.2017.11.018

[36] Desmet C, Marquette CA, Blum LJ, Doumèche B. Paper electrodes for bioelectrochemistry: biosensors and biofuel cells. Biosens Bioelectron. 2016; 76: 145–163. https://doi.org/10.1016/j.bios.2015.06.052.

[37] Mukoma P, Jooste BR, Vosloo HCM. A comparison of methanol permeability in Chitosan and Nafion 117 membranes at high to medium methanol concentrations. J Memb Sci. 2004; 243: 293–299. https://doi.org/10.1016/j.memsci.2004.06.032.

[38] Papancea A, Valente AJM, Patachia S. PVA cryogel membranes as a promising tool for the retention and separation of metal ions from aqueous solutions. J Appl Polym Sci. 2010; 118: 1567–1573. https://doi.org/10.1002/app.32514.

[39] Ma J, Sahai Y. A direct borohydride fuel cell with thin film anode and polymer hydrogel membrane. ECS Electrochem Lett. 2012; 1: F41–F43. https://doi.org/10.1149/2.005206eel.

[40] Darbari ZM, Mungray AA. Synthesis of an electrically cleanable forward osmosis membrane. Desalin Water Treat. 2016; 57: 1634–1646. https://doi.org/10.1080/19443994.2014.978390.

[41] Srinophakun P, Thanapimmetha A, Plangsri S, Vetchayakunchai S, Saisriyoot M. Application of modified chitosan membrane for microbial fuel cell: Roles of proton carrier site and positive charge. J Clean Prod. 2017; 142: 1274–1282. https://doi.org/10.1016/jjclepro.2016.06.153.

[42] Kim Y, Shin SH, Chang IS, Moon SH. Characterization of uncharged and sulfonated porous poly (vinylidene fluoride) membranes and their performance in microbial fuel cells. J Memb Sci. 2014; 463: 205–214. https://doi.org/10.1016/j.memsci.2014.03.061.

[43] Lee CH, Park HB, Lee YM, Lee RD. Importance of proton conductivity measurement in polymer electrolyte membrane for fuel cell application. Ind Eng Chem Res. 2005; 44: 7617–7626. https://doi.org/10.1021/ie0501172.

[44] Ji E, Moon H, Piao J, Ha PT, An J, Kim D, et al. Interface resistances of anion exchange membranes in microbial fuel cells with low ionic strength. Biosens Bioelectron. 2011; 26: 3266–3271. https://doi.org/10.1016/j.bios.2010.12.039.

[45] Zinadini S, Zinatizadeh AA, Rahimi M, Vatanpour V, Rahimi Z. High power generation and COD removal in a microbial fuel cell operated by a novel sulfonated PES/PES blend proton exchange membrane. Energy. 2017; 125: 427–438 https://doi.org/10.1016/j.energy.2017.02.146.

[46] Day TJ, Schmitt UW, Voth GA. The mechanism of hydrated proton transport in water. J Am Chem Soc. 2000; 122: 12027–12028. https://doi.org/10.1021/ja002506n.

[47] Mohammadifar M, Zhang J, Yazgan I, Sadik O, Choi, S. Power-on-paper: Origami-inspired fabrication of 3-D microbial fuel cells. Renew Energ. 2018; 118: 695–700. https://doi.org/10.1016/j.renene.2017.11.059

[48] Veerubhotla R, Das D, Pradhan D. A flexible and disposable battery powered by bacteria using eyeliner coated paper electrodes. Biosens Bioelectron. 2017; 94: 464–470. https://doi.org/10.1016/j.bios.2017.03.020

[49] Yee RSL, Rozendal RA, Zhang K, Ladewig BP. Cost effective cation exchange membranes: A review. Chem Eng Res Des. 2012; 90: 950–959. https://doi.org/10.1016/j.cherd.2011.10.015.

[50] Chiellini E, Corti A, D’Antone S, Solaro R. Biodegradation of poly (vinyl alcohol) based materials. Prog Polym Sci. 2003; 28: 963–1014. https://doi.org/10.1016/S0079-6700(02)00149-1.

[51] Zawodzinski TA, Neeman M, Sillerud LO, Gottesfeld S. Determination of water diffusion coefficients in perfluorosulfonate ionomeric membranes. J Phys Chem. 1991; 95: 6040–6044. https://doi.org/10.1021/j100168a060

[52] Lefebvre O, Shen Y, Tan Z, Uzabiaga A, Chang IS, Ng HY. A comparison of membranes and enrichment strategies for microbial fuel cells. Bioresour Technol. 2011; 102: 6291–6294. https://doi.org/10.1016/j.biortech.2011.02.003.

[53] Ayyaru S, Dharmalingam S. Development of MFC using sulphonated polyether ether ketone (SPEEK) membrane for electricity generation from waste water. Bioresour Technol. 2011; 102: 11167–11171. https://doi.org/10.1016/j.biortech.2011.09.021.

[54] Hernández-Flores G, Poggi-Varaldo HM, Solorza-Feria O, Ponce-Noyola MT, Romero-Castañón T, Rinderknecht-Seijas N, et al. Characteristics of a single chamber microbial fuel cell equipped with a low cost membrane. Int J Hydrogen Energy. 2015; 40 17380–17387. https://doi.org/10.1016/j.ijhydene.2015.10.024.

[55] Hernández-Flores G, Poggi-Varaldo HM, Solorza-Feria O, Romero-Castañón T, Ríos-Leal E, Galíndez-Mayer J. Batch operation of a microbial fuel cell equipped with alternative proton exchange membrane. Int. J. Hydrogen Energy. 2015; 40: 17323–17331. https://doi.org/10.1016/j.ijhydene.2015.06.057.

[56] Venkatesan PN, Dharmalingam S. Development of cation exchange resin-polymer electrolyte membranes for microbial fuel cell application. J Mater Sci. 2015; 50: 6302–6312. https://doi.org/10.1007/s10853-015-9167-x.

[57] Sivasankaran A, Sangeetha D, Ahn YH. Nanocomposite membranes based on sulfonated polystyrene ethylene butylene polystyrene (SSEBS) and sulfonated SiO_2_ for microbial fuel cell application. Chem Eng J. 2016; 289: 442–451. https://doi.org/10.1016/jxej.2015.12.095.

[58] Tiwari BR, Noori MT, Ghangrekar MM. A novel low cost polyvinyl alcohol-Nafion-borosilicate membrane separator for microbial fuel cell. Mater Chem Phys. 2016; 182: 86–93. https://doi.org/10.1016/j.matchemphys.2016.07.008.

[59] Angioni S, Millia L, Bruni G, Ravelli D, Mustarelli P, Quartarone E. Novel composite polybenzimidazole-based proton exchange membranes as efficient and sustainable separators for microbial fuel cells. J Power Sources. 2017; 348: 57–65. https://doi.org/10.1016/jjpowsour.2017.02.084.

[60] Chouler J, Bentley I, Vaz F, O’Fee A, Cameron PJ, Di Lorenzo M. Exploring the use of cost-effective membrane materials for microbial fuel cell based sensors. Electrochim Acta. 2017; 231: 319–326. https://doi.org/10.1016/j.electacta.2017.01.195.

[61] Harewood, AJT, Popuri SR, Cadogan EI, Lee CH, Wang CC. Bioelectricity generation from brewery wastewater in a microbial fuel cell using chitosan/biodegradable copolymer membrane. Int. J. Environ. Sci. Technol. 2017; 14: 1535–1550. https://doi.org/10.1007/s13762-017-1258-6.

